# Does generalism drive pollinators’ abundance, persistence and regional distribution? Insights from an ecological model

**DOI:** 10.64898/2025.12.03.692198

**Authors:** Andrea Coppola, Lorenzo Mari, Renato Casagrandi

## Abstract

The functioning of plant-pollinator mutualistic networks is crucial for ecosystem service provisioning and biodiversity maintenance. However, multiple drivers of global change are causing an alarming decline of wild pollinators’ abundance and richness. We propose an ecological, process-based mathematical model describing the dynamics of pollinators and plants, properly mediated by reward resources. Our model explicitly accounts for the main interactions of both facilitative and competitive nature that occur both within and between the two guilds. We apply our model to a broad set of real communities in fragmented landscapes to investigate the mechanisms that link the architecture of the interaction networks, the pollinators’ temporal persistence and abundance at the community level, and their rarity at the landscape level. Our results suggest that few generalist pollinators form a core of abundant, persistent and widely distributed species, while a lower number of mutualistic partners is generally associated with low abundance, low persistence and high turnover between patches. Specialists, however, are crucial to maintaining high levels of biodiversity within the community. This finding highlights the importance of ecological connectivity, through which local extinctions can be counter-balanced by recolonizations. Our analysis shows how a mechanistic model accounting for the structure of plant-pollinator networks can serve as a tool to investigate important ecological mechanisms driving community composition, dynamics and the resulting species distribution patterns.

## 1 Introduction

The worldwide decline of pollinators is an ecological issue of major concern, due to their importance for the maintenance of terrestrial biodiversity and agricultural productivity (Ollerton et al. 2011; Potts et al. 2016). Plants and pollinators within a community are intertwined by complex networks of interactions (Bascompte et al. 2007), and the architecture of these networks have been the object of a broad amount of research, carried out with both empirical and theoretical approaches.

From a theoretical point of view, the results of numerous modeling studies suggest that network architecture plays a major role on how these communities respond to global environmental change, as it can substantially affect community resilience, structural stability and species persistence (Rohr et al. 2014; Lever et al. 2014; Domínguez-Garcia et al. 2024). Testing on the field these theoretical findings on how network structure affects community dynamics is not trivial, because real-world mutualistic networks exhibit marked variability in both time and space, due to the temporal and spatial turnover of species and their set of interactions, as shown by several empirical studies (Olesen et al. 2008; CaraDonna et al. 2020; Sabatino et al. 2010; Aizen et al. 2012).

However, there is growing evidence showing how the magnitude of the observed turnover (in both time and space) is linked to the species position within the network (*i*.*e*. the topological features of the corresponding node), with interaction between generalist forming a core of ubiquitous and temporally persistent interactions, and interactions involving specialists tending to be more infrequent in both time and space (Chacoff et al. 2018; Resasco et al. 2021; Zografou et al. 2020). Moreover, empirical studies on long time series suggest that pollinators with higher specialization are the ones experiencing a more marked decline (Burkle et al. 2013; Zoller et al. 2023).

In the present study, we apply a novel process-based model to a large dataset to investigate the ecological mechanisms linking pollinators’ degree, their expected persistence and abundance in the local communities and the patch occupancy within the region.

## 2 Material and methods

### 2.1 The Dataset

We compiled data from five different studies, each of them describing a set of plant-pollinator communities scattered across a fragmented landscape. In each of these studies, the authors identified a number of patches within a region, and, for each patch, they identified the local community, in terms of both species composition and interaction network. The overall dataset comprises around 700 pollinator species and 600 plant species, organized in 85 networks, from as many patches, distributed across five different regions. References and additional information on the empirical data are available in Table 1.

**Table 1:**
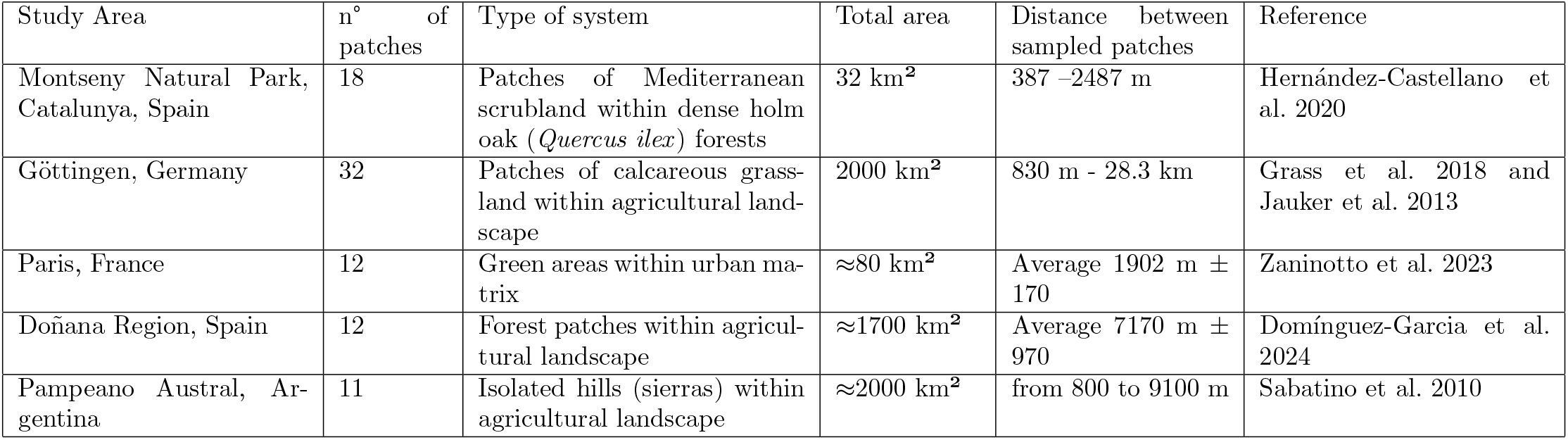
summary of study areas and their characteristics. Data are publicly available within the Supplementary Information of References in this table and/or within the Supplementary Information of Galiana et al. 2022

### 2.2 The model

#### 2.2.1 Outline and assumptions

Our model takes into account both competitive and mutualistic interactions, in order to describe population dynamics of each plant and pollinator species within a community. Flower resources produced by each plant species are also explicitly modeled as state variables. Following an approach already proposed by Bastolla et al. 2009 and Rohr et al. 2014, we assume that the benefit received by plants through pollination is simply a saturating function of the abundance of their mutualistic partners.

In turn, the benefit pollinators receive from plants is not simply modeled as a function of plants abundance, but derives from an actual resource-consumer interaction between pollinators and flower resources produced by the plants. This allows to introduce in the model an exploitative competition over common resources among pollinators that share one or more plants.

#### 2.2.2 Equations

Model equations are defined as follows:

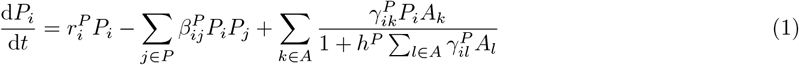

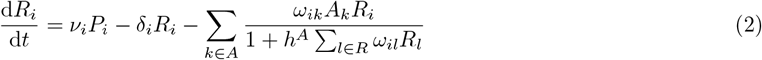

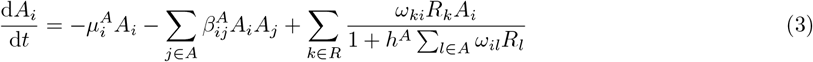

State variables P, R and A represent the abundance of plant species, their floral resources and pollinator species, respectively.

As the model works at community level, parameters concerning interactions among species can be grouped into interaction matrices. Matrix *B*^*P*^ is a square matrix whose elements are parameters describing the interspecific and intraspecific competition among plants, 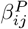 and 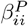, respectively. Likewise, competition among and between pollinator species is described by the square Matrix *B*^*A*^ and its elements 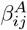 and 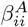. Note that, in equation 1-3, we do not impose *i* ≠*j* for nether 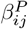 or 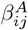.

Parameters regulating the strength of mutualistic interaction,*γ*_*ij*_ and *ω*_*ij*_, are contained matrices Γ and Ω, respectively. Matrix Γ has a number of rows and columns equal to the number of plant ad pollinator species present in the community, while the dimension of Ω match Γ’s transpose.

Parameter *γ*_*ij*_ represents the mutualistic benefit of pollinator *j* on plant *i* per unit of abundance, while parameter *ω*_*ij*_ represents the attack rate of pollinator species *j* foraging on the resource produced by plant *i*. If plant *i* and pollinator *j* do not interact, both *γ*_*ij*_ and *ω*_*ij*_ are set to zero.

The population dynamics of plant *i* (*P*_*i*_), described by equation 1, is modeled as follows: a positive intrinsic growth rate 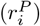 and intraspecific competition (*β*_*ii*_) leads to a logistic growth in absence of other species. When other plant species are present, they affect *P*_*i*_ dynamics via competition for soil, nutrients, sunlight and so on. Adopting the same approach of Bastolla et al. 2009 and Lever et al. 2014, in presence of *k* mutualistic partners, *P*_*i*_ receives a benefit that increases yet saturates with the partners’abundance. Parameters *γ*_*ik*_ and *h*^*P*^ defines the Holling’s type 2 functional response. Equation 2 describes the dynamics of floral resources of plant *i* (*R*_*i*_). Floral resources are produced proportionally to plant abundance, are subject to a natural decay and are consumed by the *k* pollinators that interact with plant *i*. parameter *ν*_*i*_ represent the resource production rate, parameter *δ*_*i*_ is the resource decaying rate. Resource consumption is governed by a Holling’s type 2 functional response, where *ω*_*ik*_ and *h*_*ik*_ represent attack rate and handling time of pollinator *j* consuming resource *i*, respectively.

We assume that no pollinator species is able to survive in the absence of all floral resources, therefore, in equation 3, 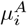 represents the per capita death rate of pollinator due to hunger mortality. A second negative term parameter accounts for the intra- and interspecific competition, shaped by the *B*^*A*^ matrix. In presence of *n* mutualistic partners, pollinator *i* receives a benefit by foraging on each of the *n* floral resources. The benefit is described, once again, by a Holling’s type II functional response, multiplied by conversion efficiency *ϵ*.

#### 2.2.3 Application to real world networks

We apply the model to all the 85 real-word networks of the dataset by running *n* = 100 simulations for each network. Before each simulation, we randomly sample all the parameters (except *ϵ, h*^*A*^ and *h*^*A*^) from gaussian distributions. We can therefore compute the persistence of each species as the fraction of simulations in which that species has an equilibrium abundance higher than a set extinction threshold (*τ* ^*A*^ and *τ* ^*P*^ for pollinators and plants, respectively).

This approach to compute persistence is based on the idea of structural stability and it is similar in concept to the one proposed by Domínguez-Garcia et al. 2024. The second metric that we computed is the average equilibrium abundance of each pollinators among the *n* simulations (from now on, simply called abundance). More info on parameter values is available in supplementary material.

For each species in each network, we also compute the eigenvalue centrality type II ( EV2, as defined by Halekotte et al. 2025). This third metric is not obtained via model simulations, it is instead based only on network topology, taking into account both the pollinator’s degree and the degree of its plant partners, and it is computed as follows:

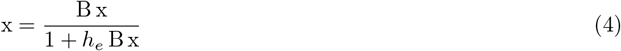

where *B* is the adjacency matrix and *h*_*e*_ is a parameter modulating saturation, here assumed equal to 1 for simplicity.

## 3 Results

Despite the geographic distance and differences in landscape type, our results show consistent patterns in both spatial distribution data and simulation outcomes. With the aim of highlighting these similarities, all the figure reported in this section are obtained aggregating across all the five regions of the dateset. Results divided by region are reported in the supplementary materials.

Figure 2 shows two patterns that emerge when grouping pollinators according to their maximum observed degree across their region. This metric, corresponding to the widest observed trophic niche, is a measure of the “level of generalism” of a certain pollinator. Grey bars shows how there is a strong heterogeneity in the maximum degree distribution, with the vast majority of pollinators that exhibit consistently a low degree in every patch. Green box-plots shows the pattern between regional distribution of species across patches and the local distribution of species within the network of interactions. In particular, pollinator with a higher maximum degree (generalists) tend to be more common ( *i*.*e*. occupy more patches) than specialists. In other words, generalists are more rare than specialists in terms of species richness, but they tend to occupy a higher number of sites.

**Figure 1.**
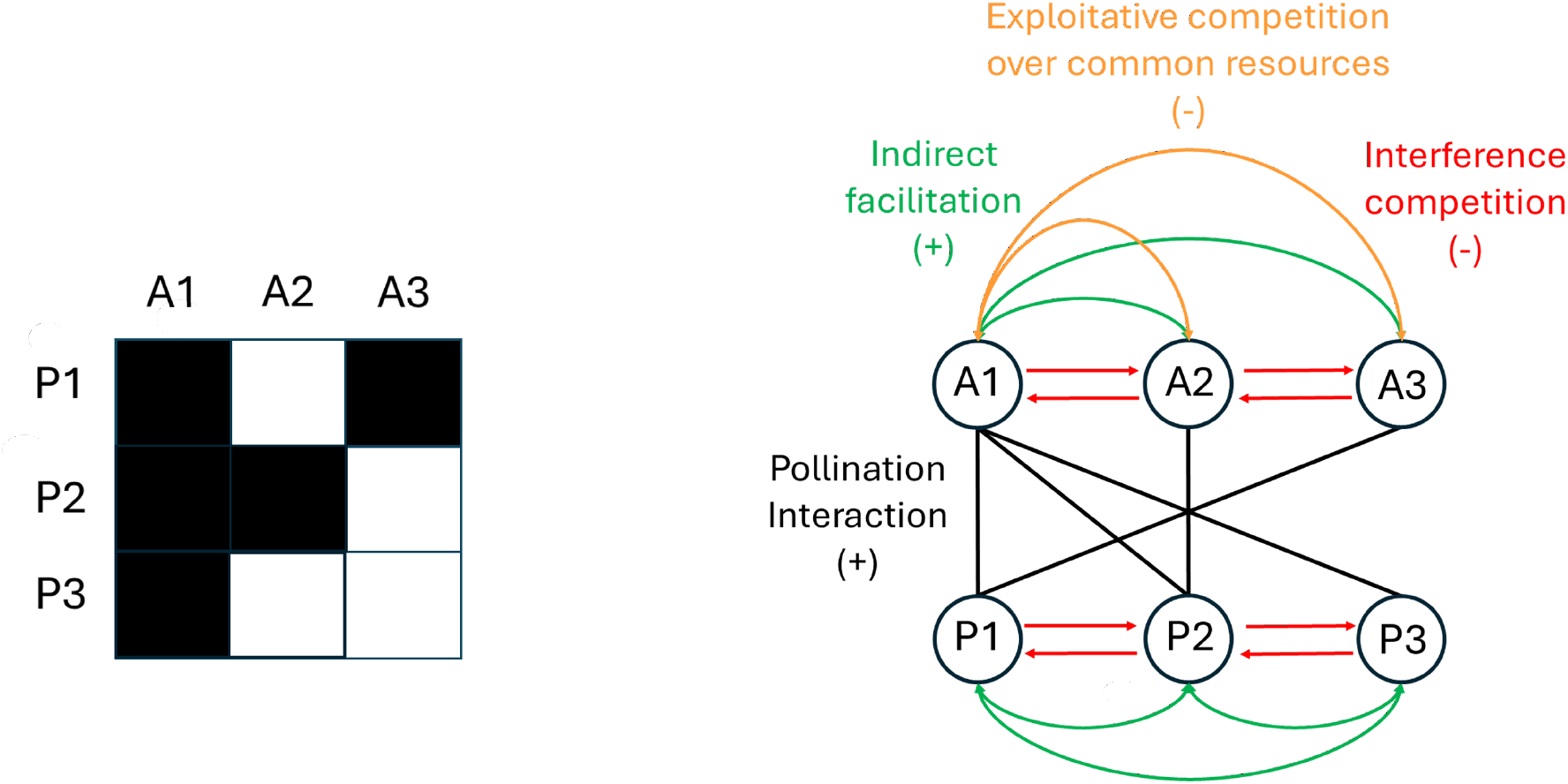
A schematic representation of the interactions included in the model, given a simple interaction matrix between three pollinator species and three plant species.

**Figure 2.**
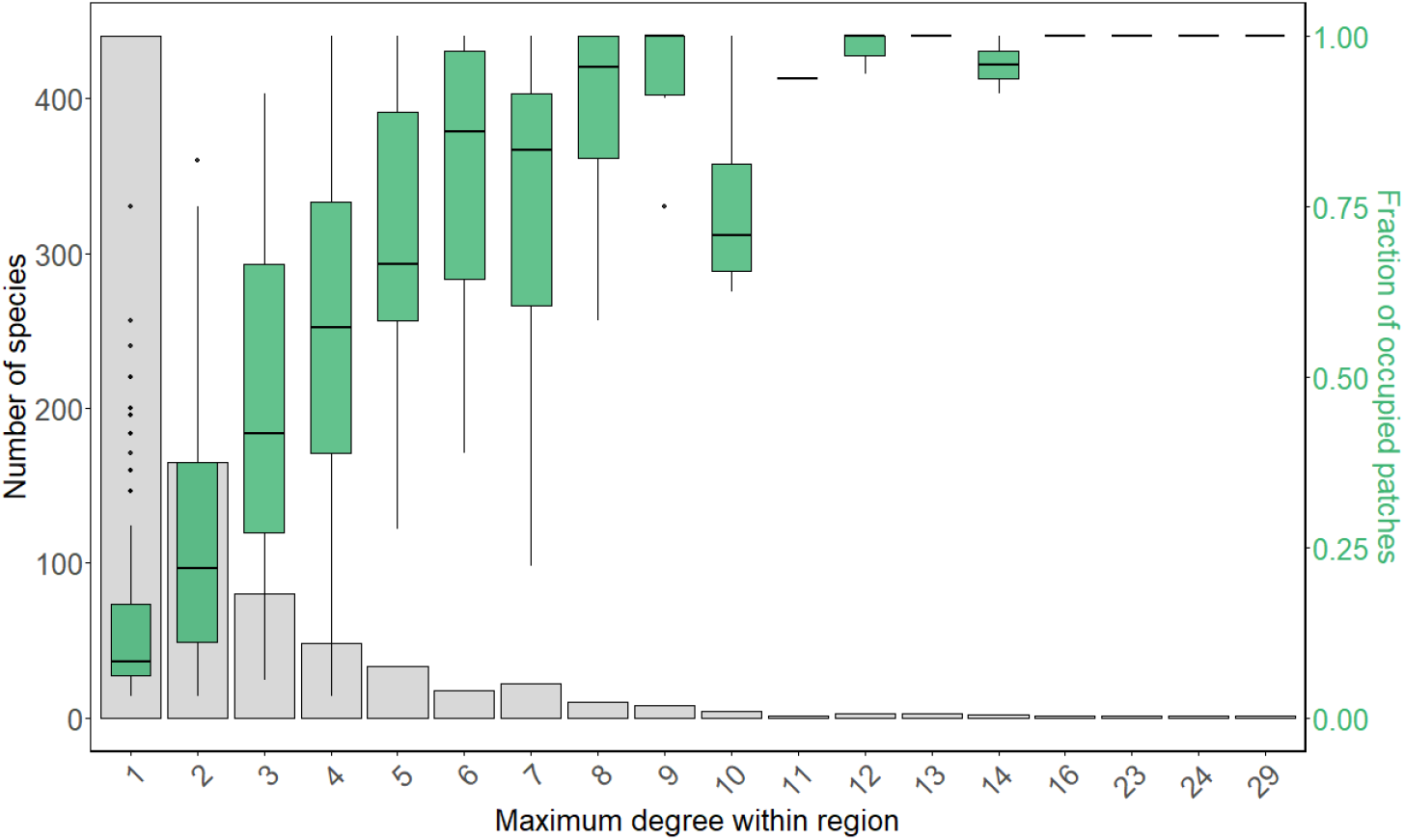
Observed distribution of number of species (gray bars) and patch occupancy (green boxplots) of pollinators when grouped by the maximum observed degree.

Concerning simulation outcomes, figure 3a shows how pollinator’s degree is positively associated with abundance and persistence, figure 3b shows that that degree is a good predictor of EV2, and figure 3c shows how within-patch performance metrics are highly correlated with EV2, thus further highlighting how a pollinator’s position within the interaction network drives its performance metrics.

**Figure 3.**
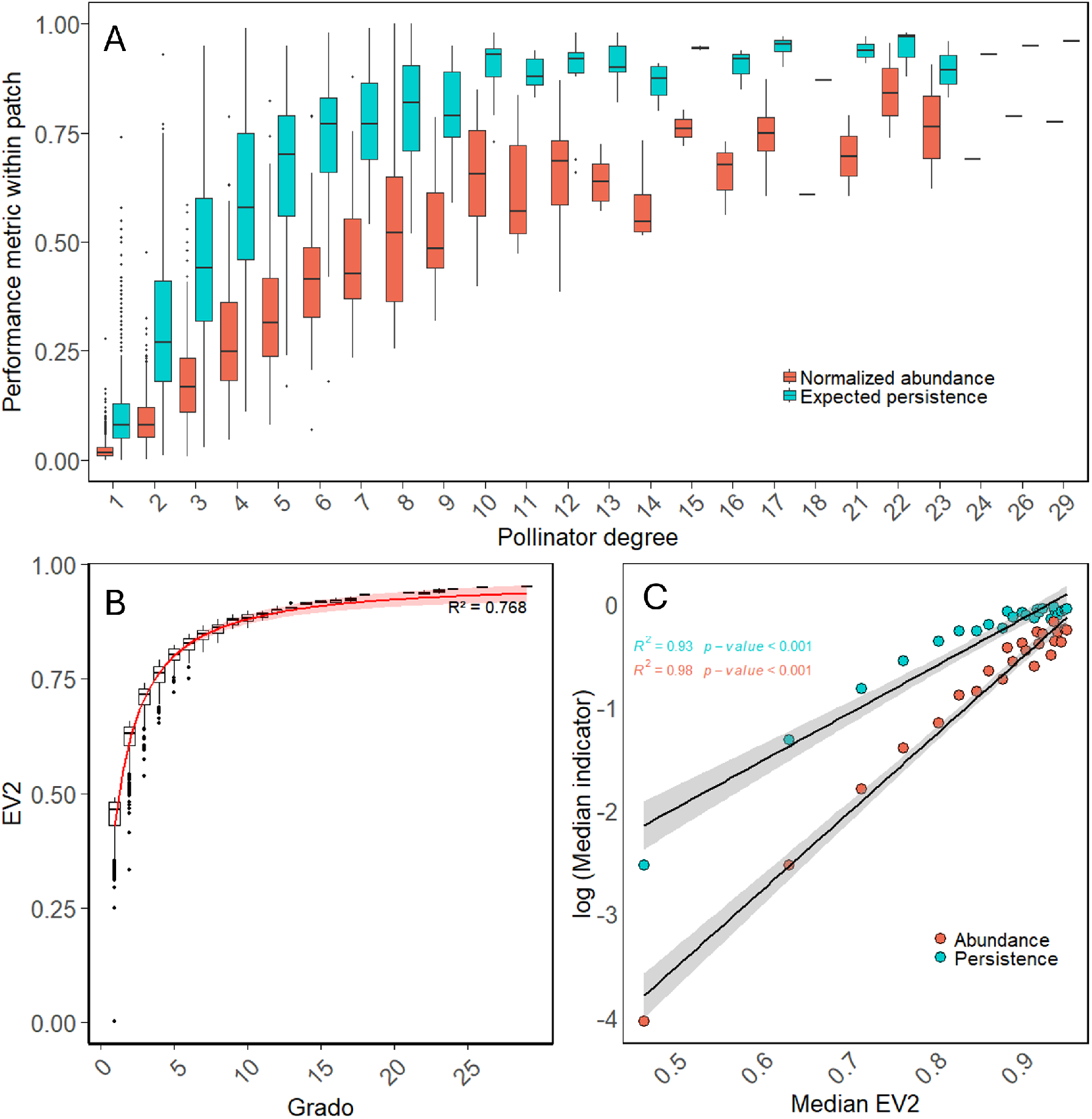
Figure 3a shows local persistence (blue boxplots) and abundance (red boxplots) of pollinators across the whole dataset, grouped by the observed degree. Figure 3b shows the correlation between observed degree and Ev2. In Figure 3C every couple of nodes vertically aligned corresponds to a degree *k*; the x-coordinate is the Median EV2 of all pollinators within the degree-*k* class, while y-coordinate is the median value of persistence (blue points) and abundance (red points) of all pollinators within that same class. Gray areas represent the 95% confidence interval of the two linear regressions, whose *R*^2^ and p-values are shown in figure.

Figure 4 contrast the observed spatial patterns with the metrics obtain via simulation of temporal dynamics.

**Figure 4.**
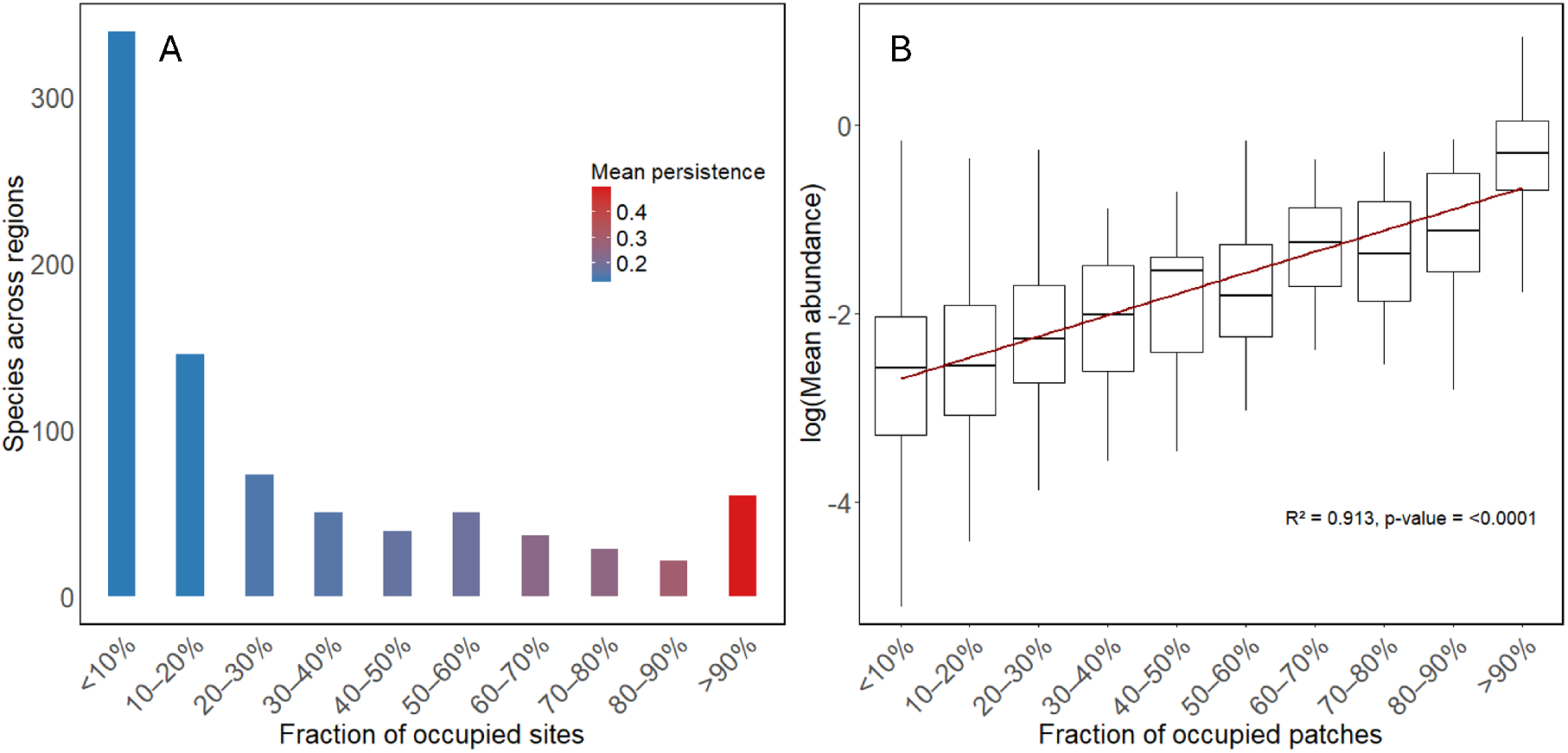
These images shows the pollinator’s occupancy frequency distribution (Figure 4a) and abundance-occupancy relationship (figure 4b).Pollinators are grouped into occupancy classes for visualization purposes. In figure 4a, the column are colored according to the mean persistence of all the pollinators falling into that class. Figure 4b is presented in semi-logarithmic scale and the regression line is computed on the median values of each boxplot.

Figure 4a shows how species number and mean persistence are distributed among occupancy classes, while Figure 4b shows instead the abundance occupancy relationship for pollinators.

The results presented here shows that pollinator communities are structured as follows: few generalist pollinators form a core of abundant, persistent and widely distributed species, while the majority of pollinators has a lower number of mutualistic partners and therefore exhibits low abundance, low persistence and high turnover between patches.

## Discussion

The relationship between generalism and abundance is at the center of a so-called “chicken egg dilemma” (Dormann et al. 2017). From one side, the abundance driven view focuses on the effects of species abundance on networks patterns, while network driven view focuses on causality in the opposite sense. Both views have been supported by empirical studies (Fort et al. 2016; Simmons et al. 2019; Song et al. 2022); However, as pointed out by Dormann et al. 2017, the two views are not mutually exclusive. As our model (and all the above mentioned ones) starts from recorded, fixed and a priori-defined interaction networks to compute community dynamics, it totally aligns with the network-driven view, thus failing in capturing the mechanisms that may lead more abundant pollinators to interact with a broader set of plants species, and vice-versa. An alternative modelling approach is represented by the work proposed by Castillo et al. 2024, where the network structure changes dynamically as new species enter into the network.

We designed the model with a minimalist approach: its equation are the simplest we could think of, with the constrain of explicitly including resource and exploitative competition among pollinators. However, its simplicity also implies the inability to include and describe several mechanism, such as interaction rewiring and adaptive foraging (see Valdovinos et al. 2013 for a more detailed and realistic model).

Despite the many inherent limitations, our model allows to propose a possible explanantion on how the set of interactions of pollinators can have a crucial role in shaping their abundance, persistence and regional distribution. More in detail, our results suggests that generalism is a driver of both local and regional abundance, while a narrow set of interactions causes the species to be both locally and regionally rare.

Halekotte et al. 2025 found high inverse correlation between the EV2 of pollinators (and plants) and their *endangerment rank*, computed leveraging a mathematical model that exhibits multiple stable states. The fact we obtain obtain a result that is qualitatively equivalent employing a different model and different performance metrics, suggest that EV2 is a reliable predictor of species’ performance in plant-pollinator networks. If this is the case, even if we found a clear link between species degree and EV2 for the 85 real networks analyzed in this study, further studies are needed to assess how much of the relationship between these two node properties is due to nestedness, core-periphery network structure or simply a high degree heterogeneity among nodes (Martín González et al. 2020; Payrató-Borràs et al. 2019).

Interestingly, pattern shown in figure 4b are coherent with the empirical data reported in Hanski 1982 and used therein as a support of the “Core-Satellite Hypothesis”. Moreover, results of further work on metapopulation models (Gyllenberg et al. 1992; Hanski et al. 1993) found that an immigration rate that increases with the fraction of occupied patches is an ecological mechanisms that can cause a positive relationship between patch occupancy and mean local abundance. The results reported in figure 4b suggest that this mechanism may not be the only one capable of generating positive abundance occupancy relationship, and that this pattern can also arise as consequence of within-patch community dynamics, driven by the local interaction networks.

On a more general note, the approach proposed in his study allows to investigate the intersection between two possible partitions of a pool of species organized in mutualistic networks. The first one is the Core-Satellite partition: proposed by Hanski 1982, concerns spatial distribution and abundance patterns, and predicts species to be either widely distributed and abundant (core species) or regionally and locally rare (satellite species). The second one is a Core-Periphery partition, that was first defined in network science (see *e*.*g*. Borgatti et al. 1999 and refernces therein) and is based on the architecture of the interaction network. This partition allows to distinguish between a core of densely connected species and a number of peripheral species, weakly connected to the network core (Morone et al. 2019; Miele et al. 2020). Our result hint towards a possible overlap between these two partitions, as pollinators that are classified as core species according to the first criteria are also core species according to the second criteria, and the same holds for satellite/peripheral species.

This hypothesis is partially as supported by empirical findings: Miele et al. 2020 studied a plant-pollinator network for six years and reported that more persistent and abundant species were more often found within the network core. However, they also showed that species position in the network was highly dynamic and that the core of species was far from being constant in time; therefore, an analysis on more empirical studies with broad spatial and temporal cover is needed to further investigate this topic.

The lack of data also strongly limits the application of mathematical models to plant-pollinator communities for forecasting, simulation under different scenarios and, perhaps in a lesser measure, system understanding. At the current state, to our knowledge, no mathematical model of plant-pollinator communities (including ours) has ever been properly calibrated and validated. Therefore, model’s output should only be considered in a qualitative way. In order to obtain more quantitatively reliable models, additional effort should be made improve the quality and the amount of data required as model-input.

First, there is a need to obtain a more comprehensive and less-biased picture of the interaction networks, complementing traditional visual observation with novel techniques such as DNA metabarcoding (Pornon et al. 2017) and automated monitoring (Serra-Marin et al. 2025).

Secondly, there is a need to move forward the soft mean field approach, where the parameters of all species of the same guild are imposed equal (or extracted from the same distribution). Modelers need data on the species traits, in order to effectively calibrate species-specific parameters, thus improving the realism of the proposed models.

Third, there is a need to gather more data describing the temporal dynamics of the communities and meta-communities, in order to validate the proposed models ( examples of long community-level time series can be found in Gaiarsa et al. 2021, Zografou et al. 2020 and Domínguez-Garcia et al. 2024).

Further insights on how species persist at landscape level could be obtained expanding the model in order to take into account i) the size of different patches, thus incorporating species-area relationship (SAR) and network area relationships (NAR) (Galiana et al. 2022) and ii) the dispersal of individuals between patches, thus up-scaling the analysis at the meta-community level (Gawecka et al. 2023).

Generalist pollinators have been indicated as of particular conversation interest, as they occur more consistently in time and space, and form a densely-connected network core that might support the more rare and inconsistent specialists (Zografou et al. 2020). Nevertheless, the vast majority of pollinators present in the communities consid-ered in this study can be classified as specialists, as they only interact with a small subset of the plant species and are characterized by low abundance, low persistence and low patch occupancy, thus implying a strong turnover, in both space and time. The survival of these species across the landscape is therefore heavenly relying on ecological connectivity, that ensures a flow of individuals between patches, thus allowing local extinctions to be countered by recolonizations. If the patches were isolated, our model predicts that only a small fraction of the sampled species would be able to persist in the long term, leading to loss of both alpha- and beta-diversity. This interpretation of the result highlights the importance of connectivity for species persistence, is coherent with the findings of several other studies (Brodie et al. 2025 and references therein) and may guide potential actions of landscape planning aimed at enhancing and preserving the biodiversity of plant-pollinator communities.

## Acknowledgments

Project funded under the National Recovery and Resilience Plan (NRRP), Mission 4 Component 2 Investment 1.4 - Call for tender No. 3138 of 16 December 2021, rectified by Decree n.3175 of 18 December 2021 of Italian Ministry of University and Research funded by the European Union – NextGenerationEU; Award Number: Project code CN 00000033, Concession Decree No. 1034 of 17 June 2022 adopted by the Italian Ministry of University and Research, D43C22001250001, Project title “National Biodiversity Future Center – NBFC”.

## References

Aizen, Marcelo A., Malena Sabatino, and Jason M. Tylianakis (Mar. 2012). “Specialization and rarity predict nonrandom loss of interactions from mutualist networks”. In: Science 335.6075, pp. 1486–1489. ISSN: 10959203. DOI: 10.1126/SCIENCE.1215320/ASSET/707DAB31-4CD9-44DA-BAF1-E0E0E5922F71/ASSETS/GRAPHIC/335_1486_F2.JPEG. URL: https://www.sciencemag.org/cgi/content/full/335/6075/1483/DC1.

Bascompte, Jordi and Pedro Jordano (2007). Plant-animal mutualistic networks: The architecture of biodiversity. DOI: 10.1146/annurev.ecolsys.38.091206.095818.

Bastolla, Ugo et al. (Apr. 2009). “The architecture of mutualistic networks minimizes competition and increases biodiversity”. In: Nature 2009 458:7241 458.7241, pp. 1018–1020. ISSN: 1476-4687. DOI: 10.1038/nature07950. URL: https://www.nature.com/articles/nature07950.

Borgatti, Stephen P and Martin G Everett (1999). Models of corerperiphery structures. Tech. rep., pp. 375–395. URL: https://www.elsevier.comrlocatersocnet.

Brodie, Jedediah F. et al. (Apr. 2025). A well-connected Earth: The science and conservation of organismal movement. DOI: 10.1126/science.adn2225.

Burkle, Laura A., John C. Marlin, and Tiffany M. Knight (Mar. 2013). “Plant-pollinator interactions over 120 years: Loss of species, co-occurrence, and function”. In: Science 340.6127, pp. 1611–1615. ISSN: 10959203. DOI: 10.1126/science.1232728.

CaraDonna, Paul J. and Nickolas M. Waser (Sept. 2020). “Temporal flexibility in the structure of plant–pollinator interaction networks”. In: Oikos 129.9, pp. 1369–1380. ISSN: 1600-0706. DOI: 10.1111/OIK.07526. URL: https://onlinelibrary.wiley.com/doi/full/10.1111/oik.07526 https://onlinelibrary.wiley.com/doi/abs/10.1111/oik.07526 https://nsojournals.onlinelibrary.wiley.com/doi/10.1111/oik.07526.

Castillo, William J., Laura A. Burkle, and Carsten F. Dormann (Dec. 2024). “Dynamics of a plant–pollinator network: extending the Bianconi–Barabási model”. In: Applied Network Science 9.1. ISSN: 23648228. DOI: 10.1007/s41109-024-00636-0.

Chacoff, Natacha P., Julian Resasco, and Diego P. Vázquez (Jan. 2018). “Interaction frequency, network position, and the temporal persistence of interactions in a plant–pollinator network”. In: Ecology 99.1, pp. 21–28. ISSN: 00129658. DOI: 10.1002/ecy.2063.

Domínguez-Garcia, Virginia et al. (2024). “Interaction network structure explains species’ temporal persistence in empirical plant–pollinator communities”. In: Nature Ecology and Evolution. ISSN: 2397334X. DOI: 10.1038/s41559-023-02314-3.

Dormann, Carsten F et al. (2017). “Identifying Causes of Patterns in Ecological Networks: Opportunities and Limitations”. In: Evolution, and Systematics Downloaded from https://www.annualreviews.org. Guest 12, p. 47. DOI: 10.1146/annurev-ecolsys-110316. URL: https://doi.org/10.1146/annurev-ecolsys-110316-.

Fort, Hugo, Diego P. Vázquez, and Boon Leong Lan (Jan. 2016). “Abundance and generalisation in mutualistic networks: Solving the chicken-and-egg dilemma”. In: Ecology Letters 19.1, pp. 4–11. ISSN: 14610248. DOI: 10.1111/ele.12535.

Gaiarsa, Marília Palumbo, Claire Kremen, and Lauren C. Ponisio (June 2021). “Pollinator interaction flexibility across scales affects patch colonization and occupancy”. In: Nature Ecology and Evolution 5.6, pp. 787–793. ISSN: 2397334X. DOI: 10.1038/s41559-021-01434-y.

Galiana, Núria et al. (Mar. 2022). “Ecological network complexity scales with area”. In: Nature Ecology and Evolution 6.3, pp. 307–314. ISSN: 2397334X. DOI: 10.1038/s41559-021-01644-4.

Gawecka, Klementyna A. and Jordi Bascompte (Aug. 2023). “Habitat restoration and the recovery of metacommunities”. In: Journal of Applied Ecology 60.8, pp. 1622–1636. ISSN: 13652664. DOI: 10.1111/1365-2664.14445.

Grass, Ingo et al. (Sept. 2018). “Past and potential future effects of habitat fragmentation on structure and stability of plant–pollinator and host–parasitoid networks”. In: Nature Ecology and Evolution 2.9, pp. 1408–1417. ISSN: 2397334X. DOI: 10.1038/s41559-018-0631-2.

Gyllenberg, Mats and Ilkka Hanski (1992). Single-Species Metapopulation Dynamics: A Structured Model. Tech. rep., pp. 35–61.

Halekotte, Lukas, Anna Vanselow, and Ulrike Feudel (Sept. 2025). “Keep the bees off the trees: the vulnerability of species in the periphery of mutualistic networks to shock perturbations”. In: Journal of Physics: Complexity 6.3. ISSN: 2632072X. DOI: 10.1088/2632-072X/ade927.

Hanski, I (1982). Dynamics of Regional Distribution: The Core and Satellite Species Hypothesis. Tech. rep. 2, pp. 210–221.

Hanski, Ilkka and Mats Gyllenbergt (1993). TWO GENERAL METAPOPULATION MODELS AND THE CORE-SATELLITE Tech. rep. 1, pp. 17–41. URL: http://www.journals.uchicago.edu/t-and-c.

Hernández-Castellano, Carlos et al. (July 2020). “A new native plant in the neighborhood: effects on plant–pollinator networks, pollination, and plant reproductive success”. In: Ecology 101.7. ISSN: 19399170. DOI: 10.1002/ecy.3046.

Jauker, Birgit et al. (Jan. 2013). “Linking life history traits to pollinator loss in fragmented calcareous grasslands”. In: Landscape Ecology 28.1, pp. 107–120. ISSN: 15729761. DOI: 10.1007/s10980-012-9820-6.

Lever, J. Jelle et al. (Mar. 2014). “The sudden collapse of pollinator communities”. In: Ecology Letters 17.3, pp. 350–359. ISSN: 1461-0248. DOI: 10.1111/ELE.12236. URL: https://onlinelibrary.wiley.com/doi/full/10.1111/ele.12236 https://onlinelibrary.wiley.com/doi/abs/10.1111/ele.12236 https://onlinelibrary.wiley.com/doi/10.1111/ele.12236.

Martín González, Ana M. et al. (Apr. 2020). Core-periphery structure in mutualistic networks: an epitaph for nestedness? DOI: 10.1101/2020.04.02.021691. URL: http://biorxiv.org/lookup/doi/10.1101/2020.04.02.021691.

Miele, Vincent, Rodrigo Ramos-Jiliberto, and Diego P. Vázquez (July 2020). “Core–periphery dynamics in a plant–pollinator network”. In: Journal of Animal Ecology 89.7, pp. 1670–1677. ISSN: 13652656. DOI: 10.1111/1365-2656.13217.

Morone, Flaviano, Gino Del Ferraro, and Hernán A. Makse (Jan. 2019). “The k-core as a predictor of structural collapse in mutualistic ecosystems”. In: Nature Physics 15.1, pp. 95–102. ISSN: 17452481. DOI: 10.1038/s41567-018-0304-8.

Olesen, Jens M. et al. (June 2008). “TEMPORAL DYNAMICS IN A POLLINATION NETWORK”. In: Ecology 89.6, pp. 1573–1582. ISSN: 1939-9170. DOI: 10.1890/07-0451.1. URL: https://onlinelibrary.wiley.com/doi/full/10.1890/07-0451.1 https://onlinelibrary.wiley.com/doi/abs/10.1890/07-0451.1 https://esajournals.onlinelibrary.wiley.com/doi/10.1890/07-0451.1.

Ollerton, Jeff, Rachael Winfree, and Sam Tarrant (Mar. 2011). “How many flowering plants are pollinated by animals?” In: Oikos 120.3, pp. 321–326. ISSN: 00301299. DOI: 10.1111/j.1600-0706.2010.18644.x.

Payrató-Borras, Claudia, Laura Hernández, and Yamir Moreno (Aug. 2019). “Breaking the Spell of Nestedness: The Entropic Origin of Nestedness in Mutualistic Systems”. In: Physical Review X 9.3. ISSN: 21603308. DOI: 10.1103/PhysRevX.9.031024.

Pornon, André et al. (Dec. 2017). “DNA metabarcoding data unveils invisible pollination networks”. In: Scientific Reports 7.1. ISSN: 20452322. DOI: 10.1038/s41598-017-16785-5.

Potts, Simon G. et al. (Dec. 2016). Safeguarding pollinators and their values to human well-being. DOI: 10.1038/nature20588.

Resasco, Julian, Natacha P. Chacoff, and Diego P. Vázquez (June 2021). “Plant–pollinator interactions between generalists persist over time and space”. In: Ecology 102.6. ISSN: 19399170. DOI: 10.1002/ecy.3359.

Rohr, Rudolf P., Serguei Saavedra, and Jordi Bascompte (July 2014). “On the structural stability of mutualistic systems”. In: Science 345.6195. ISSN: 10959203. DOI: 10.1126/SCIENCE.1253497/SUPPL_FILE/ROHR-SM.PDF. URL: https://www.science.org/doi/10.1126/science.1253497.

Sabatino, Malena, Néstor Maceira, and Marcelo A. Aizen (Sept. 2010). “Direct effects of habitat area on interaction diversity in pollination webs”. In: Ecological Applications 20.6, pp. 1491–1497. ISSN: 10510761. DOI: 10.1890/09-1626.1.

Serra-Marin, Pau Enric et al. (Oct. 2025). “Comparative assessment of automated and manual monitoring in comprehensive plant–pollinator communities”. In: Methods in Ecology and Evolution. ISSN: 2041-210X. DOI: 10.1111/2041-210X.70165. URL: https://besjournals.onlinelibrary.wiley.com/doi/10.1111/2041-210X.70165.

Simmons, Benno I. et al. (Sept. 2019). “Abundance drives broad patterns of generalisation in plant–hummingbird pollination networks”. In: Oikos 128.9, pp. 1287–1295. ISSN: 16000706. DOI: 10.1111/oik.06104.

Song, Chuliang et al. (Sept. 2022). “Generalism drives abundance: A computational causal discovery approach”. In: PLoS Computational Biology 18.9. ISSN: 15537358. DOI: 10.1371/journal.pcbi.1010302.

Valdovinos, Fernanda S. et al. (June 2013). “Adaptive foraging allows the maintenance of biodiversity of pollination networks”. In: Oikos 122.6, pp. 907–917. ISSN: 1600-0706. DOI: 10.1111/J.1600-0706.2012.20830.X. URL: https://onlinelibrary.wiley.com/doi/full/10.1111/j.1600-0706.2012.20830.x https://onlinelibrary.wiley.com/doi/abs/10.1111/j.1600-0706.2012.20830.x https://nsojournals.onlinelibrary.wiley.com/doi/10.1111/j.1600-0706.2012.20830.x.

Zaninotto, Vincent, Elisa Thebault, and Isabelle Dajoz (Feb. 2023). “Native and exotic plants play different roles in urban pollination networks across seasons”. In: Oecologia 201.2, pp. 525–536. ISSN: 14321939. DOI: 10.1007/s00442-023-05324-x.

Zografou, Konstantina et al. (Aug. 2020). “Stable generalist species anchor a dynamic pollination network”. In: Ecosphere 11.8. ISSN: 21508925. DOI: 10.1002/ecs2.3225.

Zoller, Leana, Joanne Bennett, and Tiffany M. Knight (Jan. 2023). “Plant–pollinator network change across a century in the subarctic”. In: Nature Ecology and Evolution 7.1, pp. 102–112. ISSN: 2397334X. DOI: 10.1038/s41559-022-01928-3.

