## Supplementary information for "Does generalism drive pollinators’ abundance, persistence and regional distribution? Insights from an ecological model"

[1] Dipartimento di Elettronica, Informazione e Bioingegneria, Politecnico di Milano, 20133 Milano, Italy.

### Supplementary materials

#### Results per Region

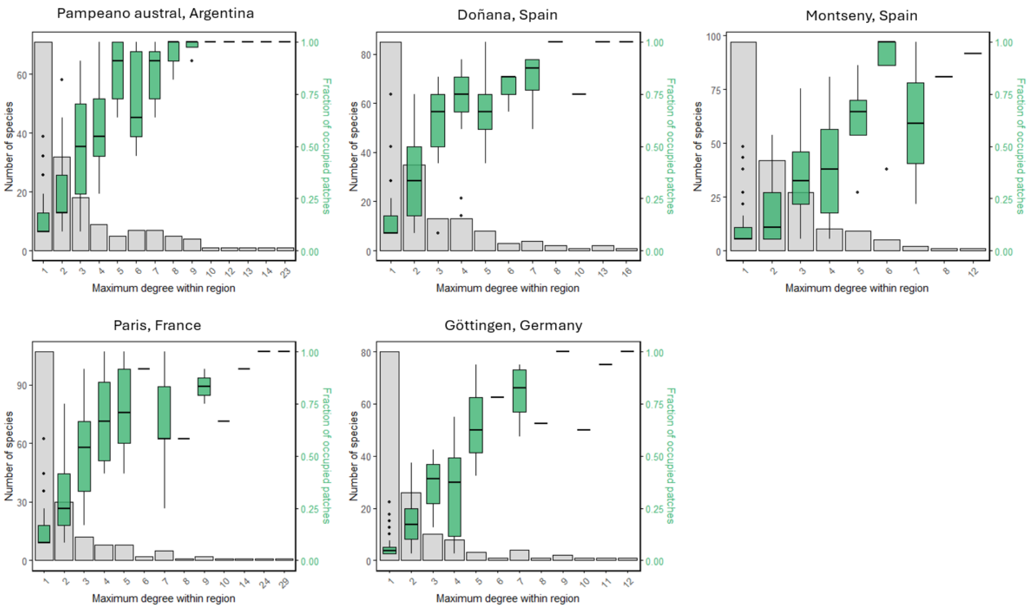


Figure 1: Observed distribution of number of species (gray bars) and patch occupancy (green boxplots) of pollinators when grouped by the maximum observed degree.

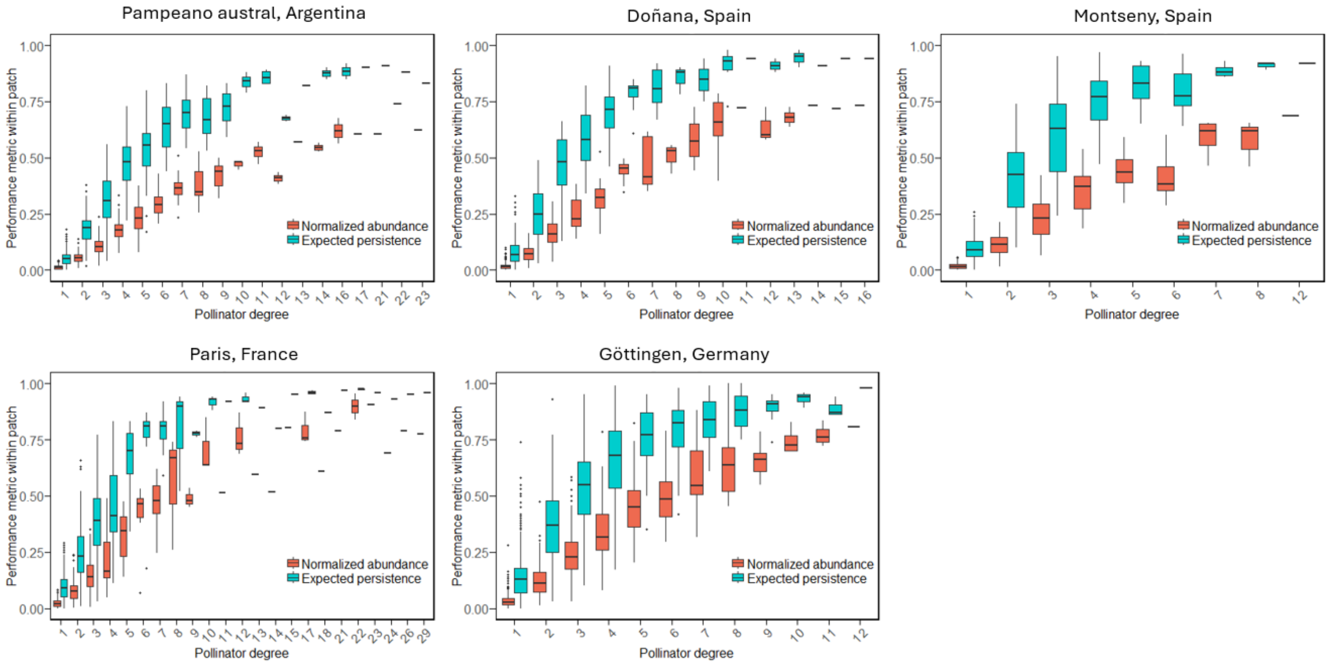


Figure 2: Figure S3 shows local persistence (blue boxplots) and abundance (red boxplots) of pollinators across the whole dataset, grouped by the observed degree.

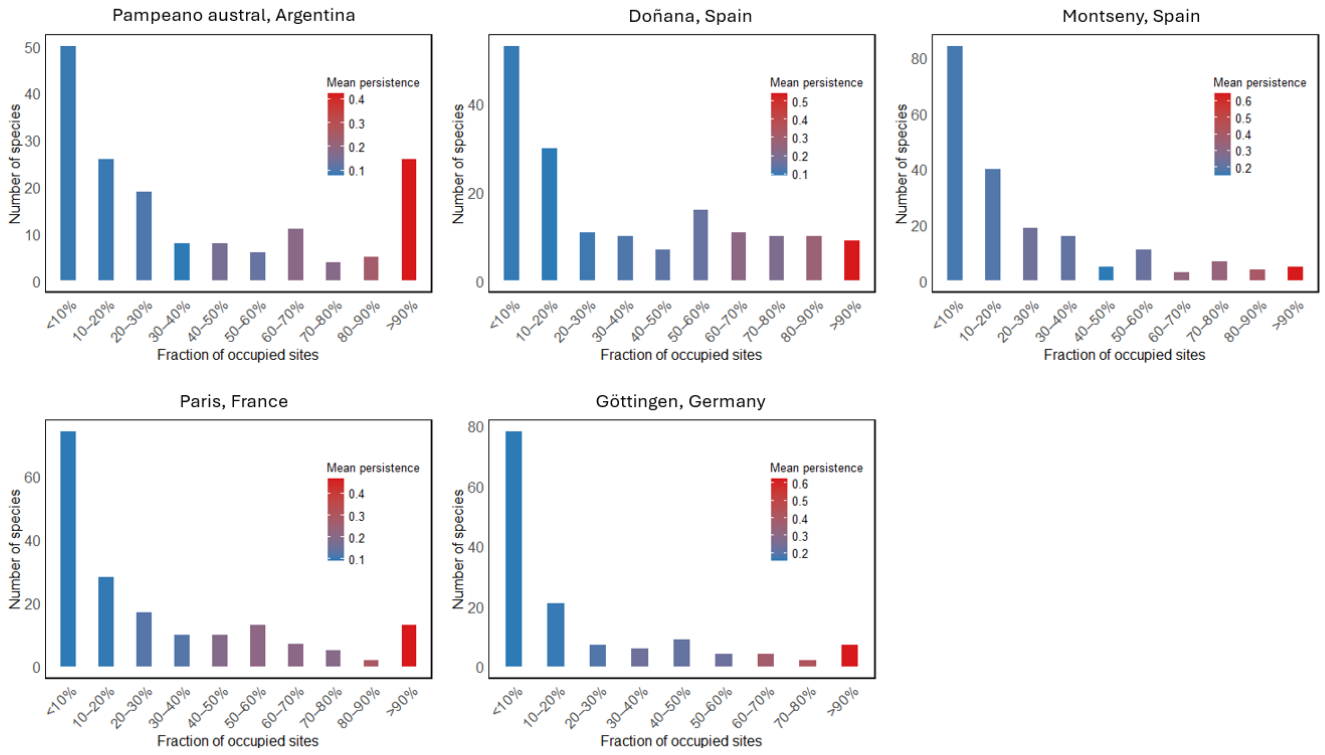


Figure 3: Pollinator's occupancy frequency distribution, columns are colored according to the mean persistence of all the pollinators falling into that class.Pollinators are grouped into occupancy classes for visualization purposes.

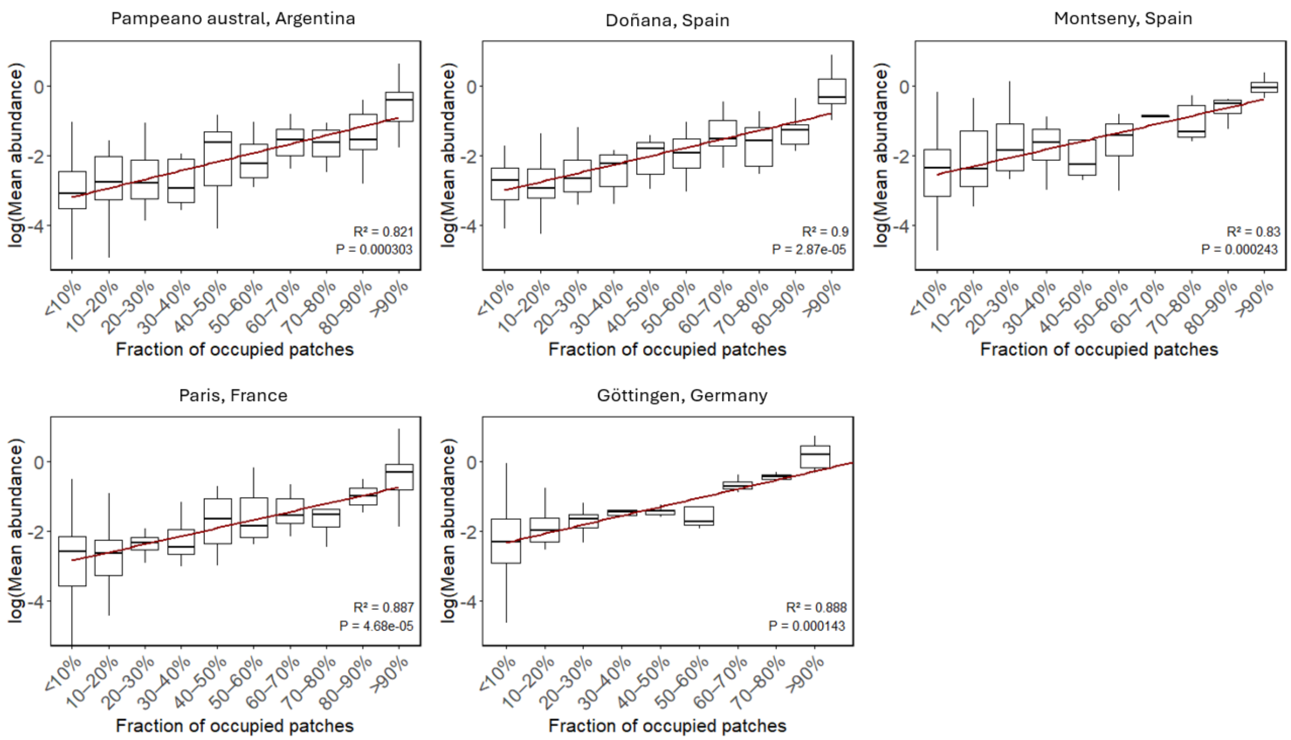


Figure 4: Pollinators Abundance-occupancy relationship. Graphs are in semi-logarithmic scale and the regression line is computed on the median values of each boxplot. Pollinators are grouped into occupancy classes for visualization purposes.

### Model parameters

| Parameter | Mean | Standard deviation | description |
|---|---|---|---|
| $r^P$ | 0.05 | 0.01 | growth rates of plants |
| $\mu^A$ | 0.1 | 0.01 | mortality rates of pollinators |
| $\beta^P$ | 0.003 | 0.005 | competition coefficients of plants |
| $\beta^A$ | 0.002 | 0.005 | competition coefficients of pollinators |
| $\gamma$ | 0.2 | 0.01 | mutualistic benefit received by plants per unit of abundance of pollinators |
| $\omega$ | 0.4 | 0.01 | attack rate of pollinators |
| $\nu$ | 0.2 | 0.01 | production rates of resources |
| $\delta$ | 0.1 | 0.001 | decay rates of resources |
| $h_P$ | 0.5 | 0 (fixed value) | plants "handling time" |
| $h_A$ | 0.5 | 0 (fixed value) | pollinators handling time |
| $\epsilon$ | 0.1 | 0 (fixed value) | conversion efficiency of pollinators |
| $\tau^A$ | 0.1 | 0 (fixed value) | extinction threshold for pollinators |
| $\tau^P$ | 0.1 | 0 (fixed value) | extinction threshold for pollinators |
